# 3-D structure based prediction of SUMOylation sites

**DOI:** 10.1101/2022.08.19.504594

**Authors:** Yogendra Ramtirtha, M. S. Madhusudhan

**Affiliations:** Indian institute of Science Education and Research, Pune, India

**Keywords:** protein 3-D structures, novel, SUMOylation, sampling method

## Abstract

SUMOylation is a posttranslational modification that involves lysine residues from eukaryotic proteins. Misregulation of the modification has been linked to neuro-degenrative diseases and cancer. All the presently available tools to predict SUMOylation sites are sequence based. Here, we propose a novel structure based prediction tool to discriminate between SUMOylated and non-SUMOylated lysines. We demonstrate the method by carrying out a proof-of-concept study. The method achieved an accuracy of 81% and a Matthews’ Correlation Coefficient of 0.4.

## 1 Introduction

Lysine residues in eukaryotic proteins are known to undergo many post-translational modifications. Examples include ubiquitination, acetylation, methylation, SUMOylation etc. SUMOylation involves formation of a covalent bond between side chain amino group of lysine residues in target / substrate proteins and C-terminus of a protein called SUMO (Small Ubiquitin-related MOdifier). Candidate lysines from target proteins are selected by SUMO E2 conjugating enzyme (ubc9) independently *in vitro* or with the help of SUMO E3 ligases *in vivo*. Mutation of lysine to an arginine disrupts the modification. Disruption of SUMOylation has been linked to neuro-degenerative diseases and cancer.

Experimental determination of SUMOylated lysines is cumbersome. Hence, computational prediction of SUMOylated lysines could be useful. All the currently available SUMOylation site prediction tools make use of protein sequences. GPS-SUMO [1,2] and JASSA [3] are two such popular sequence-based SUMOylation site prediction tools. These sequence-based SUMOylation site prediction tools preferentially look for the consensus motif ψ–K-x-(E/D), where ψ– I/L/V and K – SUMOylated lysine. Recent mass spectrometry-coupled proteomics experiments conducted on human cell lines have shown that around 50% of all SUMOylated lysines conform to consensus motif [4,5]. Thus, protein sequence information alone is insufficient to predict all SUMOylated lysines. Information about protein three dimensional (3-D) structures could be useful to understand how ubc9 discriminates between SUMOylated and non-SUMOylated lysines. To the best of our knowledge, none of the currently available SUMOylation site prediction tools make direct use of protein 3-D information. Hence, this research article describes proof-of-concept study of a novel method that takes protein 3-D information into account while predicting SUMOylated lysines.

Experimental techniques such as x-ray crystallography, nuclear magnetic resonance and electron microscopy are used to solve protein structures. Protein 3-D structures are deposited in a data archive called Protein Data Bank (PDB) [6] under different accession identifiers. Structural information about known ubc9-target protein complexes is very important for designing a robust prediction method. Our current understanding of ubc9-target complexes is limited because the PDB contains information about only one ubc9-target complex where the target protein is RanGAP1 (PDB ID : 1Z5S) [7]. Given below is the image of the enzyme-target complex generated using UCSF Chimera version 1.13.1 [8] (Figure-1). Ubc9 active site has 2 important residues C93 and D127 that catalyze the formation of covalent bond between lysine side chain and SUMO C-terminal tail. Lysine binding site in ubc9 is so narrow that replacing lysine in the target protein with an arginine disrupts the formation of covalent bond.

**Figure 1:**
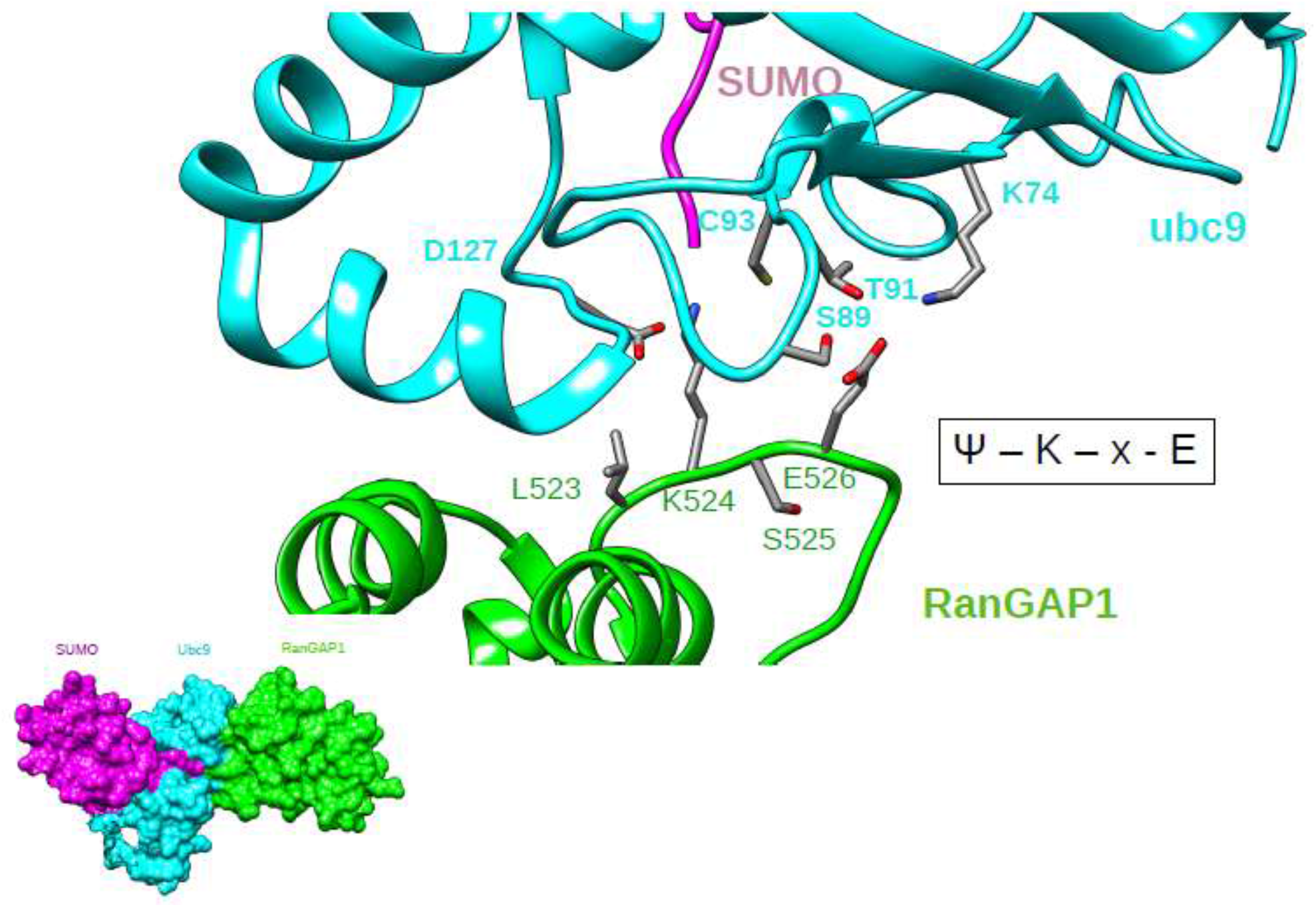
Top : Ribbon representation of a complex between SUMO (magenta), ubc9 (cyan) and RanGAP1 (green). All interacting residues are shown in ball and stick models. inset : surface representation of the same complex and same coloring scheme. SUMOylated lysine from RanGAP1 conforms to the consensus motif.

Scarcity of structural information about ubc9-target complexes is the major limiting factor for the present study. Hence, this study has been divided into three broad steps. First step involves creating a dataset of protein 3-D structures for SUMOylated proteins identified by recent experiments. All the proteins considered in this study are encoded by humans. Second step involves docking target protein structure onto ubc9 structure such that there is a lysine near the active site of ubc9. Different conformational poses of ubc9-target complex are sampled and the pose with maximum number of interprotein atomic contacts at a distance of 4Ẳ between the two proteins is chosen. Care was taken to make sure that the chosen pose did not have any atomic clashes between the main chain atoms of both the proteins. The sampling method was applied to every lysine from all the structures of the dataset and optimal pose of the ubc9-target complex obtained for every lysine was chosen. The third step involves developing a scoring method that can discriminate between ubc9-target complexes of SUMOylated and non-SUMOylated lysine. Performance of the scoring method will also be assessed/

## 2 Materials and methods

### 2.1 Generation of a dataset of SUMOylated protein structures

The list of SUMOylated proteins for this study was obtained from a recent mass-spectrometry based proteomics experiments conducted on human cell lines [5]. This list consists of 9330 proteins containing 49850 SUMOylated lysines in total. The mappings between human SUMOylated proteins and their respective Protein Data Bank (PDB) [6] identifiers were obtained with the help of SIFTS database [9,10]. Around 2331 of the 9330 SUMOylated proteins have at least one structure in the PDB containing at least one SUMOylated lysine. In order to remove redundancy in these proteins, h-CD-HIT server (http://weizhong-lab.ucsd.edu/cdhit_suite/cgi-bin/index.cgi?cmd=h-cd-hit) [11–14]was used with 3 hierarchical identity cutoffs 90%, 60% and 30% respectively. The results obtained at 30% redundancy from h-CD-HIT server contained a list of 1841 structures corresponding to 1841 SUMOylated proteins. The details of dataset used in this study are given below (Table-1). Some protein structures contained unnatural amino acids such as seleno-methionine, phospho-serine, phospho-threonine, phospho-tyrosine etc. These unnatural amino acids would have created errors during the sampling method discussed later. Hence, all the unnatural amino acids were converted to their nearest analogues from the 20 standard amino acids such as methionine, serine, threonine, tyrosine etc. Protein structures often have missing atoms because some parts of the structure have poor resolution. All such missing atoms were fixed using complete_pdb() function in MODELLER version 9.17 [15]. In some cases, residue numbering differs between UniProt sequence of a protein and the sequence of its corresponding PDB structure. In order to obtain correct residue positions, pairwise alignments were built between UniProt and PDB sequences of each protein. All pairwise alignments were built using SALIGN from MODELLER version 9.17. There are around 7432 SUMOylated lysines in the 1841 structures used in this study. All the remaining 27874 lysines in these 1841 protein structures (except the 7432 lysines) were treated as non-SUMOylated lysines.

**Table-1:** Overview of dataset used in this study

| Description | Numbers |
| --- | --- |
| Total number of target proteins | 1841 |
| Total number of SUMOylated lysines | 7432 |
| Lysines conforming to either K-x-(E/D) or (E/D)-x-K motif | 2556 |
| Lysines not conforming to consensus motif | 4876 |

### 2.2 Computational tools used in this study

All the steps in this work including dataset compilation, sampling method and scoring analysis were implemented in Python version 2.7.5. Mathematical calculations were carried out using Numeric Python (NumPy) version 1.7.1 [16]. Graphs were plotted using ggplot2 library [17] in R version 3.4.4 [18].

### 2.3 Sampling method to dock target proteins onto ubc9

It is important to understand the structure of a lysine residue before discussing the sampling method. A lysine residue has 4 main chain atoms N, CA, C and O as well as 5 side chain atoms CB, CG, CD, CE and NZ (Figure-2). The lysine main chain has 2 torsion angles phi and psi whereas its side chain has 4 torsion angles chi1, chi2, chi3 and chi4 (Figure-2). Angles between 4 atoms connected by 3 consecutive bonds are known as torsion angles.

**Figure 2:**
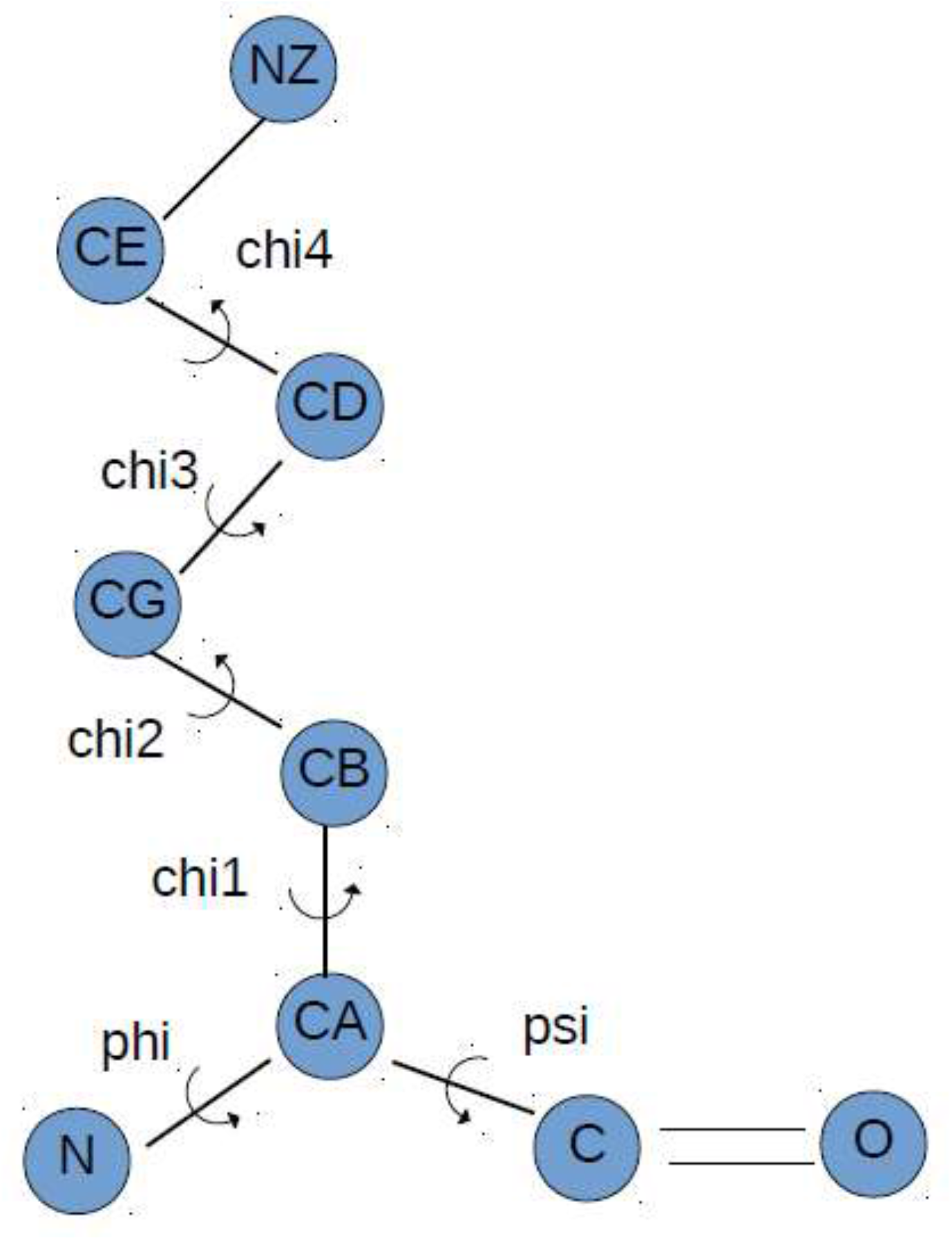
Two dimensional structure of a lysine residue.

Details of atoms involved in different torsion angles are given above (Table-2). For example, chi4 angle measures rotation of atoms around CD-CE bond and involves atoms CG, CD, CE and NZ. C_prev_ and N_next_ imply main chain C and N atoms of previous and next residues in the protein sequence respectively.

**Table-2:** Overview of atoms involved in different torsion angles

| Torsion angle | Atoms involved | Bond of interest |
| --- | --- | --- |
| Phi | C<sub>prev</sub>-N-CA-C | N-CA |
| Psi | N-CA-C-N<sub>next</sub> | CA-C |
| chi1 | N-CA-CB-CG | CA-CB |
| chi2 | CA-CB-CG-CD | CB-CG |
| chi3 | CB-CG-CD-CE | CG-CD |
| chi4 | CG-CD-CE-NZ | CD-CE |

Torsion angle calculations are important for the sampling method. These calculations depend on unit vector calculations. For a vector v = (x, y, z), unit vectors are calculated in 2 steps. First, modulus of v is calculated by taking the square root of a dot product of v with itself (Equation-1). Second, v is divided by the modulus to obtain a unit vector of v (Equation-2). In Python, dot product is calculated using the function numpy.dot() and square root is calculated using numpy.sqrt() function.

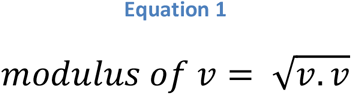

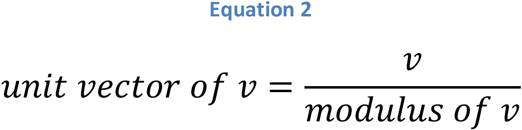

In order to understand torsion angle calculations, let us consider the chi4 angle. The chi4 angle measures rotation of atoms around CD-CE bond and it measures angle between two planes. The first plane is formed by CG, CD and CE atoms whereas the second plane is formed by CD, CE and NZ atoms. In case the four atoms have the Cartesian coordinates – CG = (x1, y1, z1), CD = (x2, y2, z2), CE = (x3, y3, z3) and NZ = (x4, y4, z4), then the torsion angle calculations go as follows:

First, we calculate vectors: b1 = (x1 – x2, y1 – y2, z1 – z2), b2 = (x3 – x2, y3 – y2, z3 – z2) and b3 = (x4 – x3, y4 – y3, z4 – z3). Second, v1, v2 and v3 are unit vectors along b1, b2 and b3 respectively. Third, vectors u1 and u3 are calculated (Equations-3 and 4).

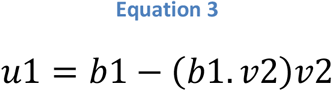

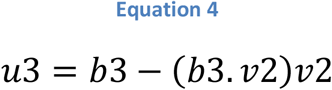

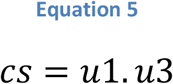

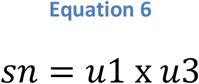

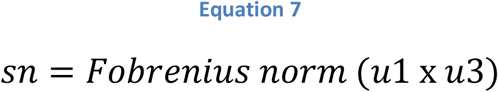

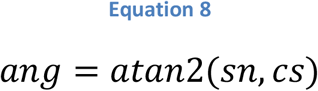

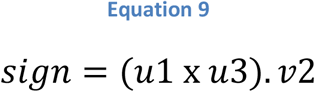

In all the above equations note the difference between dot product (.) and cross product (x). Fobrenius norm for a matrix is obtained by taking the square root of the sum of squares of all elements of the matrix. In Python, Fobrenius norm is calculated by using the function numpy.linalg.norm() and atan2 is calculated by using the function numpy.arctan2. The term “ang” represents the value of the torsion angle in radians. If the sign term is negative, then ang is also negative and hence must be multiplied by -1. The torsion angle “ang” ranges between –π to π in radians or -180° to 180°.In order to convert “ang” from radians to degrees, multiply “ang” by a factor of 180/π. Conversely, in order to convert an angle from degrees to radians, it should be multiplied by a factor of π/180.

Given above are mathematical equations necessary for torsion angle calculations (Equations-1 to 9). Now we will be discussing the mathematical equations necessary for calculating Cartesian coordinates of a rigid body after rotation. For example, let us consider 3 points – P1 = (x1, y1, z1), P2 = (x2, y2, z2) and Q = (x3, y3, z3). Axis of rotation passes through points P1 and P2, and we want to calculate Cartesian coordinates of Q after rotation by an angle θ in radians around the axis of rotation. Vector b1 = (x2 – x1, y2 – y1, z2 – z1) and u = (x0, y0, z0) is a unit vector along the direction of vector b1.

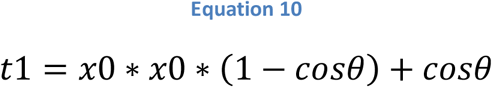

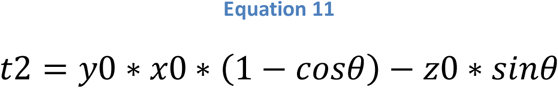

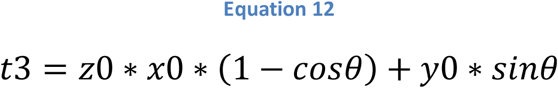

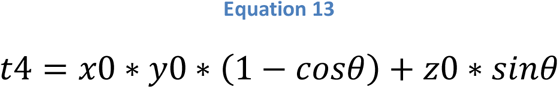

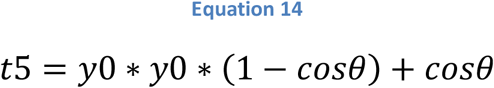

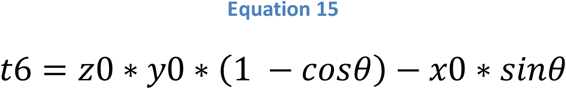

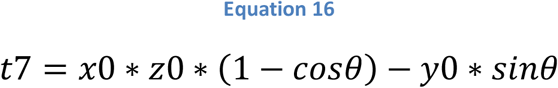

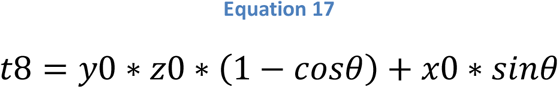

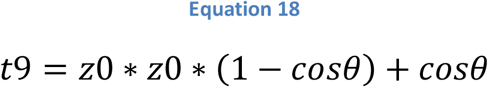

Now, we calculate vector b2 = (x4, y4, z4), where x4 = x3 – x1, y4 = y3 – y1 and z4 = z3 – z1. Let us say that (x5, y5, z50 are Cartesian coordinates of point Q after the rotational motion. The values of x5, y5 and z5 can be calculated as follows.

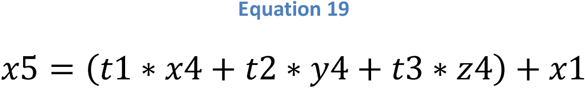

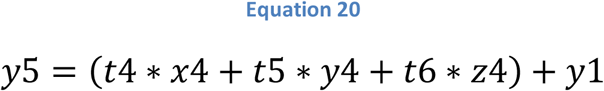

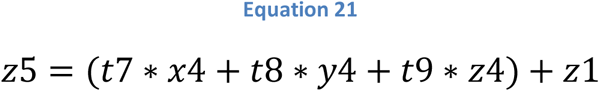

In Python, the values of sinθ and cosθ are calculated using the functions numpy.sin() and numpy.cos(). And, the value of π is obtained using the function numpy.pi(). Thus mathematical equations given above help us calculate the Cartesian coordinates of a target protein after spinning around an axis of rotation (Equations-10 to 21).

The objective of the sampling method is to dock target proteins onto ubc9 such that the lysine of interest from the target is near the active site residues of the enzyme. After docking the target onto ubc9, the method samples favorable conformational poses between the target and the enzyme. In the present method, ubc9 and target proteins are treated as rigid bodies. During the sampling process, ubc9 remains fixed whereas the target protein undergoes motions such as translations and rotations. The sampling method is carried out independently for every lysine from each of the 1841 target proteins of the dataset.

Translation is a motion wherein every atom in a rigid body is displaced by the same distance through 3-D space. Rotation is a motion wherein every atom in a rigid body spins around an axis of rotation. Schematic depiction of steps involved in the sampling method is given above (Figure-3). Each step of the sampling method is elaborated below.

**Figure 3:**
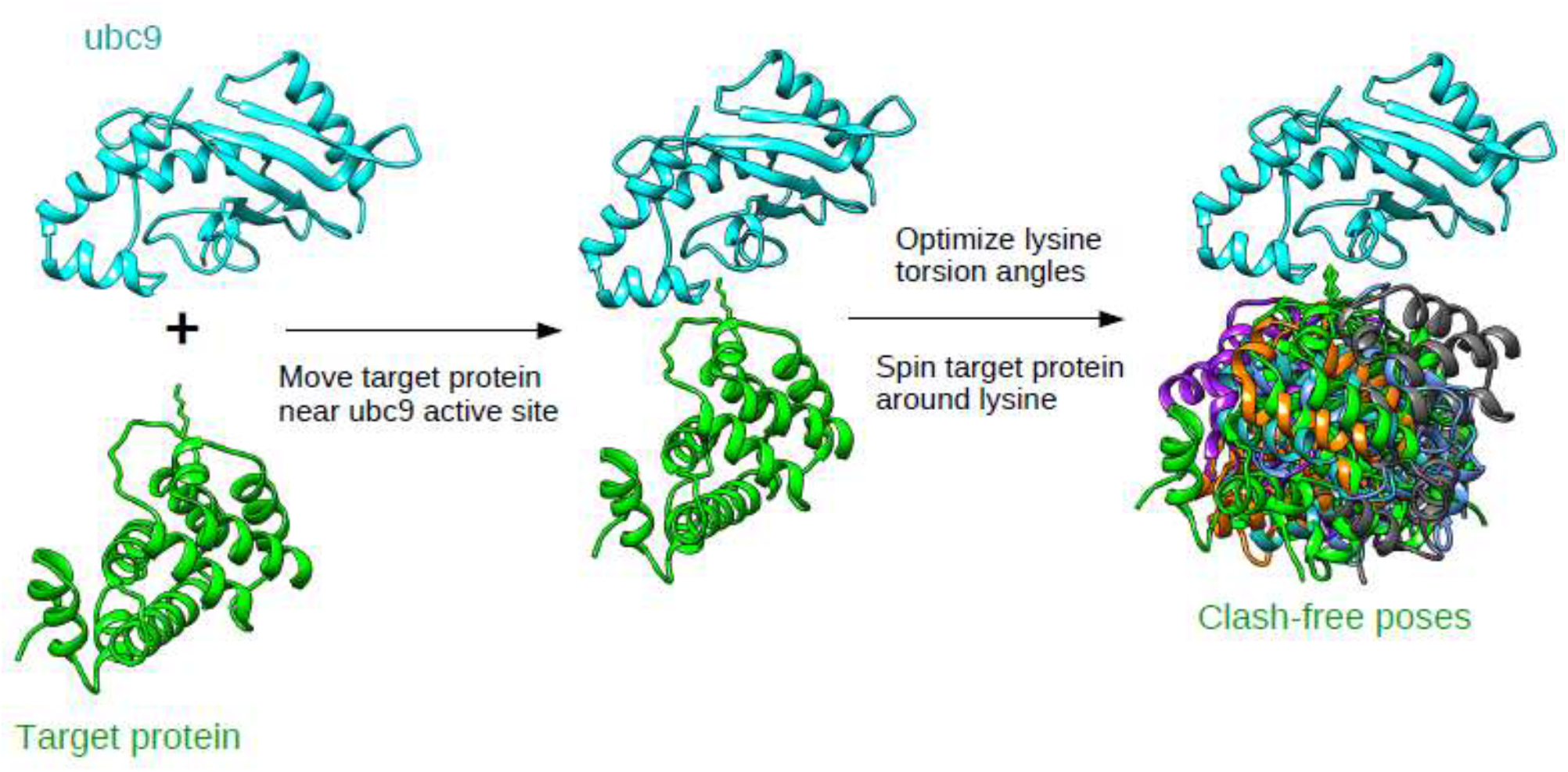
Schematic overview of the steps involved in sampling method.

#### 2.3.1 Move target protein near ubc9 active site

The first step of the sampling method aims at bringing the target protein in the vicinity of ubc9 such that the lysine of interest from the target is in the active site of the enzyme. This is achieved with the help of a technique called rigid body transformation in 3-D space, also known as 3-D least squares fit. Given below is a pictorial overview of steps involved in rigid body transformation in 3-D space (Figure 3).

A detailed explanation of rigid body transformation can be found here (http://nghiaho.com/?page_id=671) [19]. The structure of protein RanGAP1 bound to ubc9 (PDB ID : 1Z5S and Figure 1) was used as a reference for the transformation. CD, CE and NZ atoms from lysine-524 in RanGAP1 and lysine of interest in target protein are important for the transformation process. The transformation process minimizes the root mean squared deviation between both the atom sets.

Before the beginning of the transformation, the target and RanGAP1 proteins could have any arbitrary location in 3-D space (Figure 3A). Let us say, CD1 = (x1, y1, z1), CE1 = (x2, y2, z2) and NZ1 = (x3, y3, z3) are side chain atoms of lysine of interest from target protein. And, CD2 = (x4, y4, z4), CE2 = x5, y5, z5) and NZ2 = (x6, y6, z6) are side chain atoms of lysine 524 from RanGAP1. The first step of the transformation process is to translate the target protein such that the center of masses of both the above mentioned atom sets superimpose (transition of target protein from Figure 3A to 3B).

Given below are mathematical equations important for the transformation. Here, cm1 and cm2 are center of masses of both the atom sets described above. Average Cartesian coordinates are 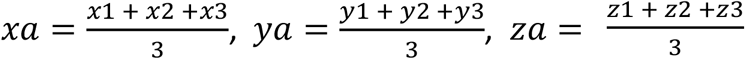 and 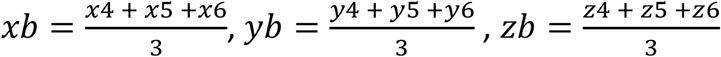.

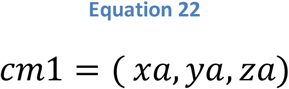

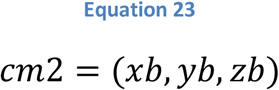

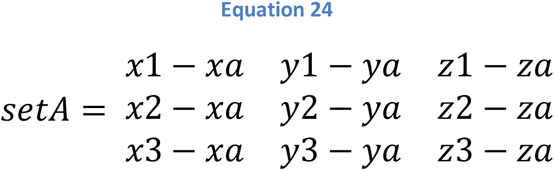

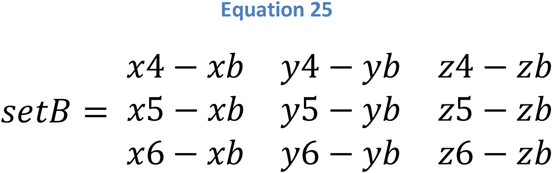

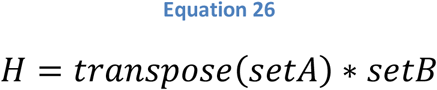

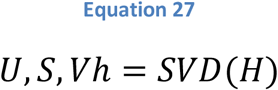

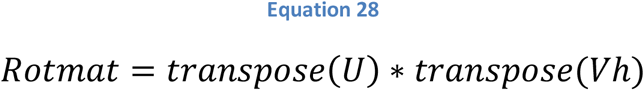

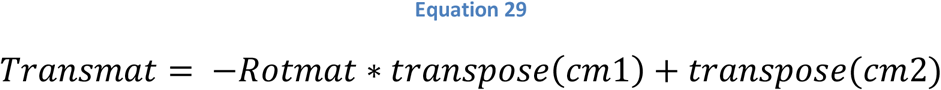

All the equations given above (Equations 22 to 29) involve matrix multiplications. “Rotmat” and “Transmat” are rotation and translation matrices that help us in calculating the Cartesian coordinates of a target protein after rigid body transformation. SVD stands for singular value decomposition (implemented using Python function numpy.linalg.svd()). After the transformation is completed, the Cartesian coordinates of RanGAP1 were deleted leaving behind the ubc9-target complex.

Rigid body transformations help in docking the target protein onto ubc9 (Figure 3A, 3B and 3C). However, the structure of the enzyme-target complex generated from transformation may not be the energetically favorable pose for the two proteins to interact. Hence, the enzyme-target complex was subjected to further conformational sampling, the details of which are given below.

#### 2.3.2 Optimize lysine torsion angles

The chi2, chi3 and chi4 torsion angles of lysine of interest from the ubc9-target complex generated above were adjusted to 172.8°, 173.8° and -175.3° respectively. This was done because the tunnel in ubc9 through which lysine accesses the active site is very narrow and accommodates lysine only in its stretched or extended conformation (Figure-1). Mutagenesis studies have shown that if lysine is substituted by arginine, then the arginine cannot enter the tunnel owing to its branched side chain. In order to adjust torsion angles to their appropriate values, all the atoms of target protein (except lysine side chain atoms) are rotated around an axis of rotation defined by bond of interest. For example, while adjusting the chi4 angle, all the atoms of target protein are rotated around CD-CE bond (Table-3) except CD, CE and NZ atoms (Table-3) of lysine side chain. Similar procedure is applied to chi1, chi2 and chi3 angles.

**Table-3:** Lysine side chain torsion angles, bond of interest and atoms kept fixed during angle adjustment

| **Torsion angle** | **Bond of interest** | **Atoms kept fixed during adjustment** |
| --- | --- | --- |
| chi1 | CA-CB | CA, CB, CG, CD, CE, NZ |
| chi2 | CB-CG | CB, CG, CD, CE, NZ |
| chi3 | CG-CD | CG, CD, CE, NZ |
| chi4 | CD-CE | CD, CE, NZ |

The list of possible values of chi1 angle was obtained from 2010 back-bone dependent rotamer library [20]. First main chain torsion angles phi and psi are calculated for the lysine of interest. Second, the rotamer library is searched for all possible lysine conformations having the given phi and psi angles as well as having average ch2, chi3 and chi4 angle values within 180 ± 10°. This list of chi1 angle values usually consists of 3 approximate conformations: -60° (+gauche), 60° (-gauche) and 180° (trans) respectively. The lysine of interest from the target protein is sampled in all the 3 conformations.

#### 2.3.3. Spin target protein around lysine

For every chi1 angle value, the target protein is subjected to an additional rotation. The axis of rotation for this motion is defined by CB and NZ atoms of lysine of interest. The target protein is spun in steps of 10°. Thus, for every chi1 angle value, there are 360° / 10° = 36 different conformational poses between ubc9 and target protein. At the end of the sampling process, a total of 3 * 36 = 108 conformational poses are sampled between ubc9 and target protein for a given lysine of interest.

Out of all the conformational poses sampled between ubc9 and target protein for a given lysine of interest, only those poses are retained that have no clashes between main chain atoms of ubc9 and target protein. Here, N, CA, C, O and CB atoms of both the proteins are considered as main chain atoms. The distance between ubc9 main chain atom A = (x1, y1, z1) and target protein main chain atom B = (x2, y2, 2z) is calculated as follows.

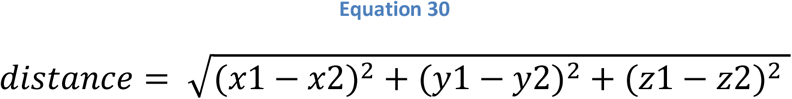

The Lennard-Jones potential (LJ) between atoms A and B can be calculated as follows.

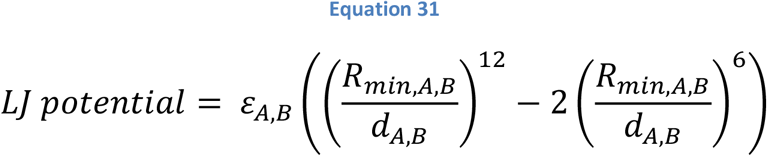

Here, *ε*_A,B_ and R_min,A,B_ are terms specific to atoms A and B that were obtained from topology parameters of AMBER 99 force field [21]. The R_min,A,B_ term is the sum of van der Waals radii of atoms A and B. When LJ potential between atoms A and B is equal to zero, Equation 31 reduces to = 0.89 * R_min,A,B_. Thus, atoms A and b are considered to be clashing if the distance between them is less than 0.89 times the sum of their van der Waals radii. At the end of sampling method, only those poses are retained that do not have clashes between main chain atoms of ubc9 and target protein. In case, a lysine of interest has more than one clash free poses, then the pose having maximum number of atomic contacts within a distance of 4Ẳ between ubc9 and target protein was chosen. This was done to ensure that one representative pose was chosen for every lysine of interest.

### 2.4 Discriminating poses based on residue contacts

The aim of this exercise is to find a combination of residue contacts that occur more in ubc9-target poses of SUMOylated lysines than in poses of non-SUMOylated lysines. This was achieved with the help of a modified version of Apriori algorithm [22]. Residue i from ubc9 and residue j from a target protein was considered to be in contact if any atom from I is within a distance of 4Ẳ from any atom of residue j. Residue contact information from all the ubc9-target poses is encoded as “res-pairs”. An example of res-pair encoding for residue contacts from ubc9-RanGAP1 complex (Figure-1) involving glutamate residues of consensus motif from RanGAP1 is given below (Table-4). Residue contacts involving lysine of interest are ignored from all ubc9-target complexes because they do not provide any new information.

**Table-4:** Example of res-pair encodings for residue contacts between ubc9 and glutamate residue of consensus motif from RanGAP1

| Ubc9 residue number : ubc9 residue type – RanGAP1 residue number : RanGAP1 residue type | Res-pair encoding |
| --- | --- |
| 89 : SER – 526 : GLU | 89 – GLU |
| 91 : THR – 526 : GLU | 91 – GLU |
| 74 : LYS – 526 : GLU | 74 – GLU |

The Apriori algorithm is commonly used for finding patterns in customer transaction data in the retail industry. Apriori algorithm clusters different items bought by customers into sets according to their support (also known as probability). For the present exercise, a res-pair could be considered as an item and res-pairs were clustered into different res-pair sets on the basis of their occurrences. The definition of support was modified as follows (Equation 32). The modified version of support was referred to as enrichment.

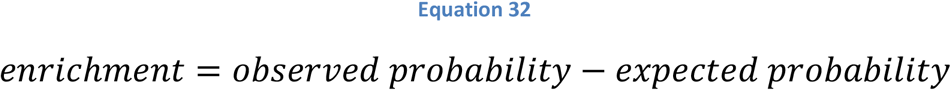

Here, observed probability refers to normalized frequency of a res-pair (or a set of res-pairs) in ubc9-target poses derived from SUMOylated lysines. Expected probability refers to normalized frequency of res-pair (or set of res-pairs) in ubc9-target poses derived from non-SUMOylated lysines. Normalization was done with respect to total number of ubc9-target poses for the given lysine category – SUMOylated or non-SUMOylated.

The Apriori algorithm was applied to 360 res-pairs that occur in ubc9-target poses of both SUMOylated and non-SUMOylated lysines. These 360 res-pairs also had absolute frequency of 40 or higher in poses of SUMOylated lysines. The Apriori algorithm can be summarized in the steps given below –

i. Calculate enrichment values for each of the 360 res-pairs. Each res-pair could be thought of as a res-pair set of size equal to 1. All res-pairs having their enrichments greater than or equal to -1.0 were selected. (There were no res-pairs having enrichments greater than or equal to 0.0).
ii. New res-pair sets were generated by extending res-pair sets from previous step by another res-pair, such that the newly added res-pair was a member of a res-pair set from previous step. All possible res-pair sets of size equal to 2 were generated.
iii. Enrichment values for all newly generated res-pair sets were calculated. All res-pair sets having enrichments greater than or equal to 0.0 were selected.
iv. This process of generation and selection of new res-pair sets having size greater by 1 than previous step is continued till no new res-pair sets can be created. At this step, the algorithm ends.

### 2.5 Statistical parameters to assess predictions

Predictions were assessed using statistical parameters such as sensitivity, specificity, accuracy, F1 score and MCC (Matthew’s Correlation Coefficient) (Equations-33 to 38). Here, TP = number of true positives, TN = number of true negatives, FP = number of false positives and FN = number of false negatives.

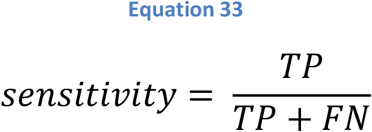

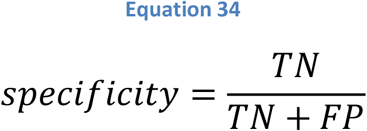

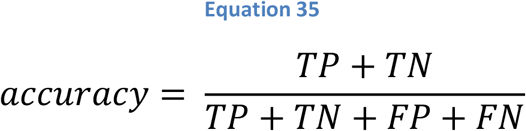

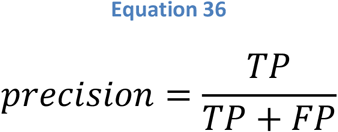

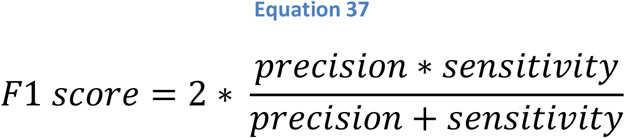

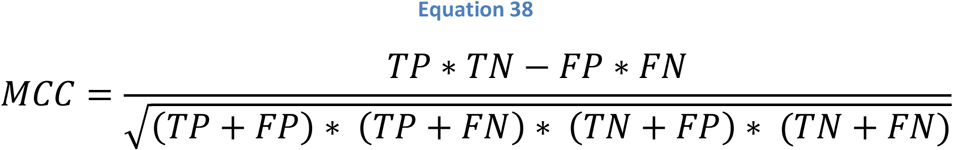

## 3 Results

### 3.1 Analysis of the target protein structures in the dataset

Majority (1349) of the protein structures used in this study were solved using x-ray crystallography (Table-5). However, a few of the structures were also solved using NMR (178), electron microscopy (313) and solution scattering (1).

**Table-5:** Experimental sources of the protein structures used in this study

| Experiment type | Number of structures |
| --- | --- |
| X-ray crystallography | 1349 |
| NMR | 178 |
| Electron microscopy | 313 |
| Solution scattering | 1 |

**Table-6:**
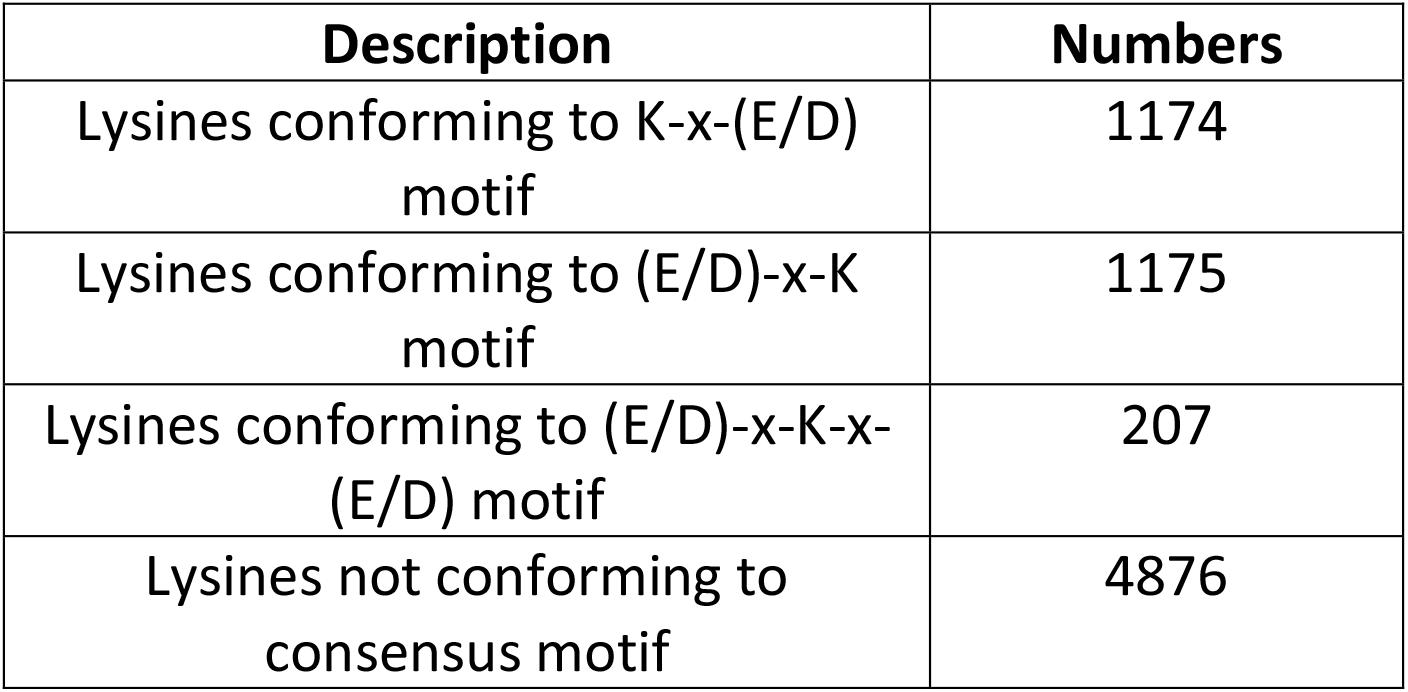
Proportion of SUMOylated lysines that conform to consensus motif

| Description | Numbers |
| --- | --- |
| Lysines conforming to K-x-(E/D) motif | 1174 |
| Lysines conforming to (E/D)-x-K motif | 1175 |
| Lysines conforming to (E/D)-x-K-x-(E/D) motif | 207 |
| Lysines not conforming to consensus motif | 4876 |

The protein structures used in this study vary in their sequence lengths (Figure 5B) as well as the number of SUMOylated lysines (Figure 5A) present in them. Most of the proteins used in this study contain 5 or less SUMOylated lysines. However, there are a handful of proteins that contain as many as 20 or more modified lysines. Similarly, majority of protein structures used in this study have sizes less than 1000 amino acids. There are a handful of structures that have sizes equal to or greater than 2000 amino acids. Thus, SUMOylation occurs in proteins of varying sizes and varying number of lysines in these proteins.

**Figure 4:**
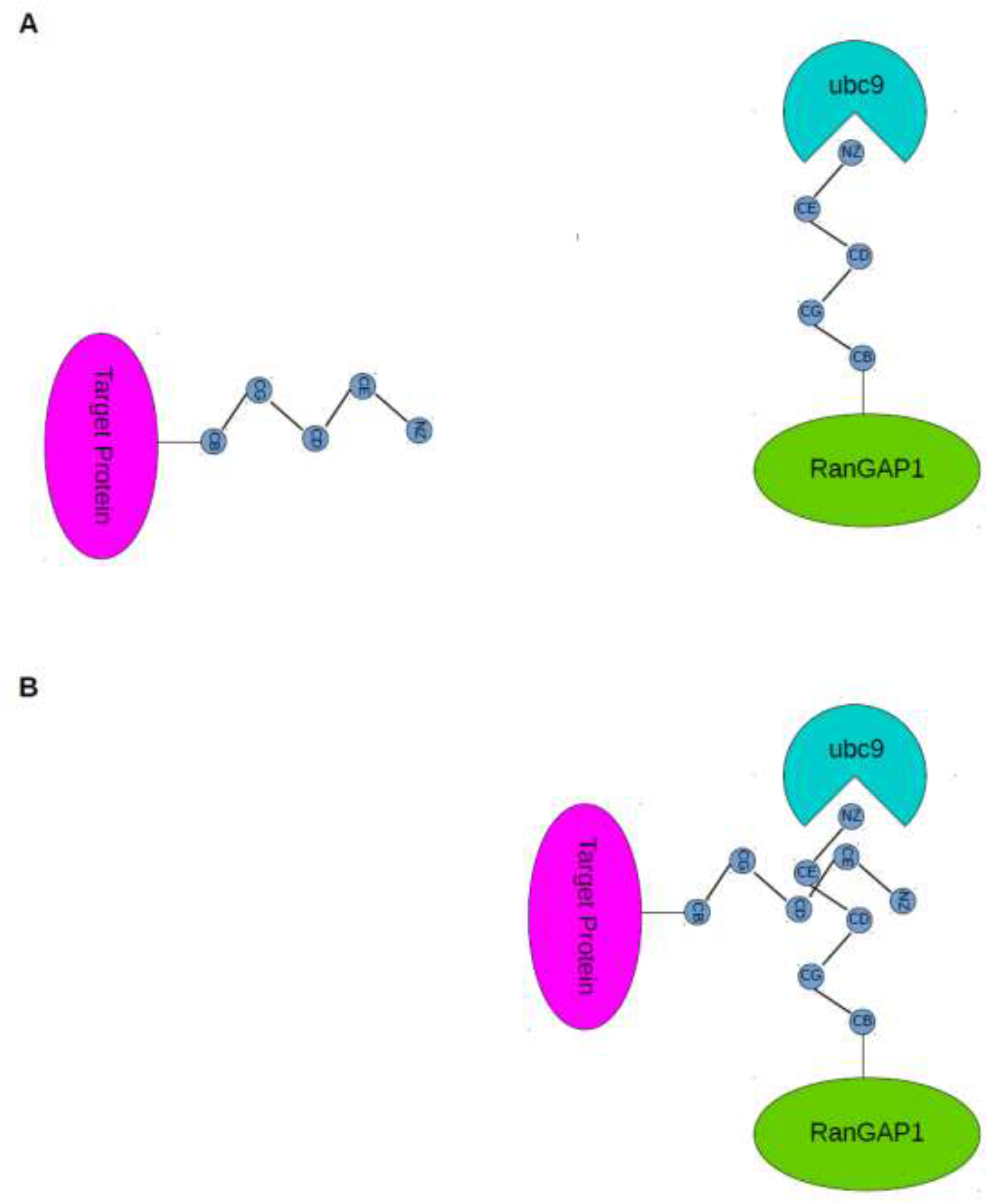
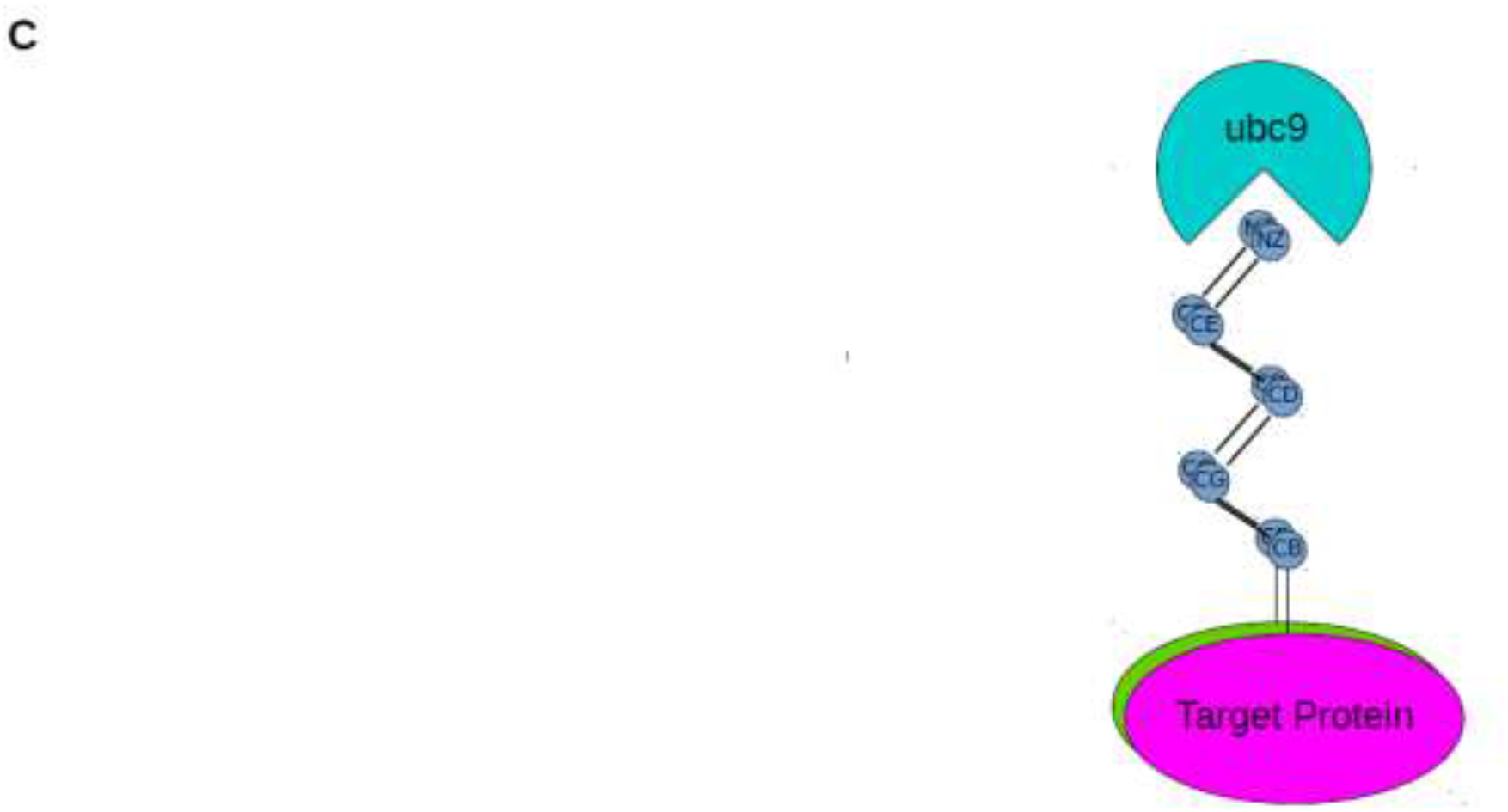
Schematic representation of steps involved in rigid body transformation of target protein in 3-D space.

**Figure 5:**
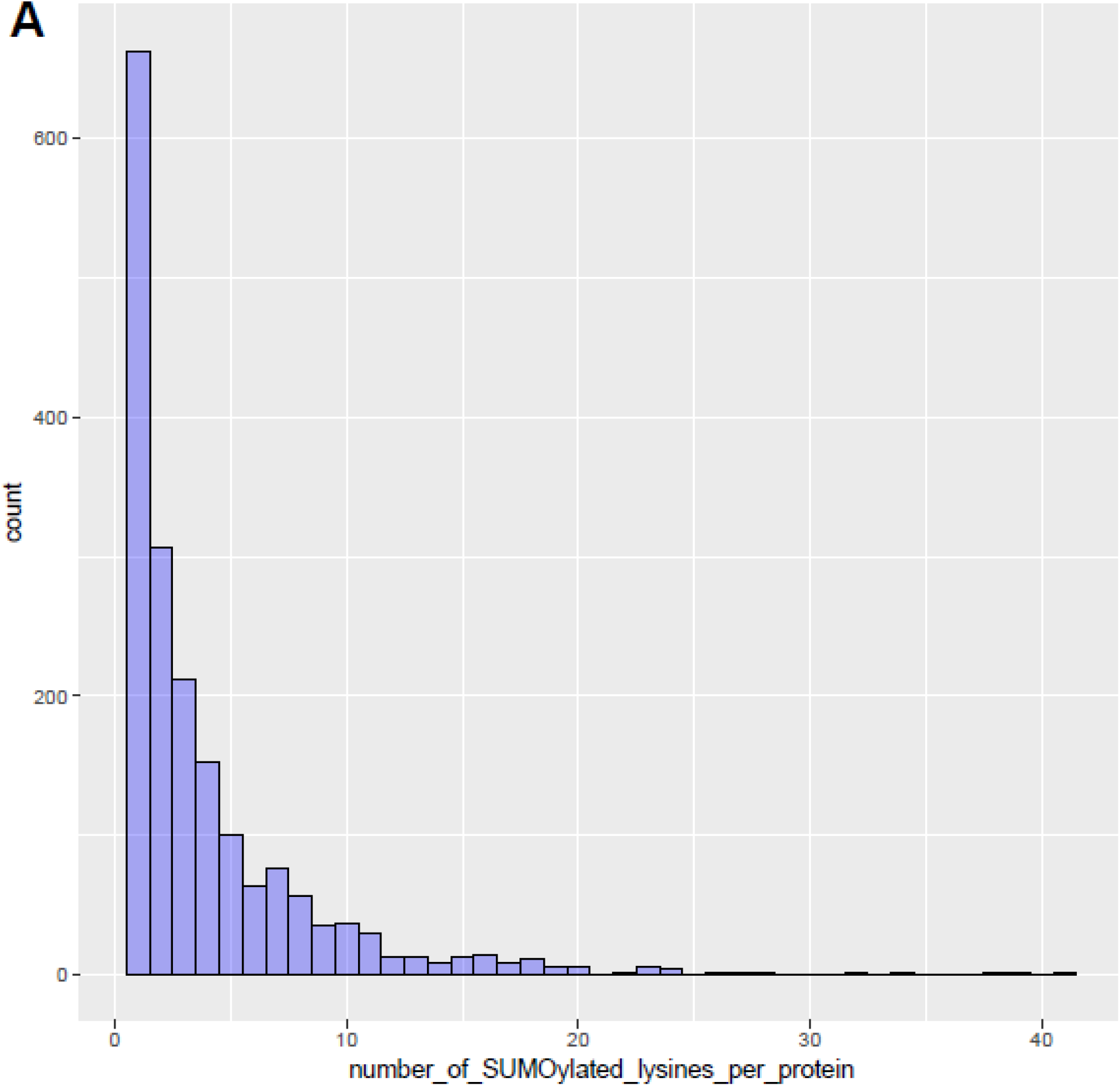
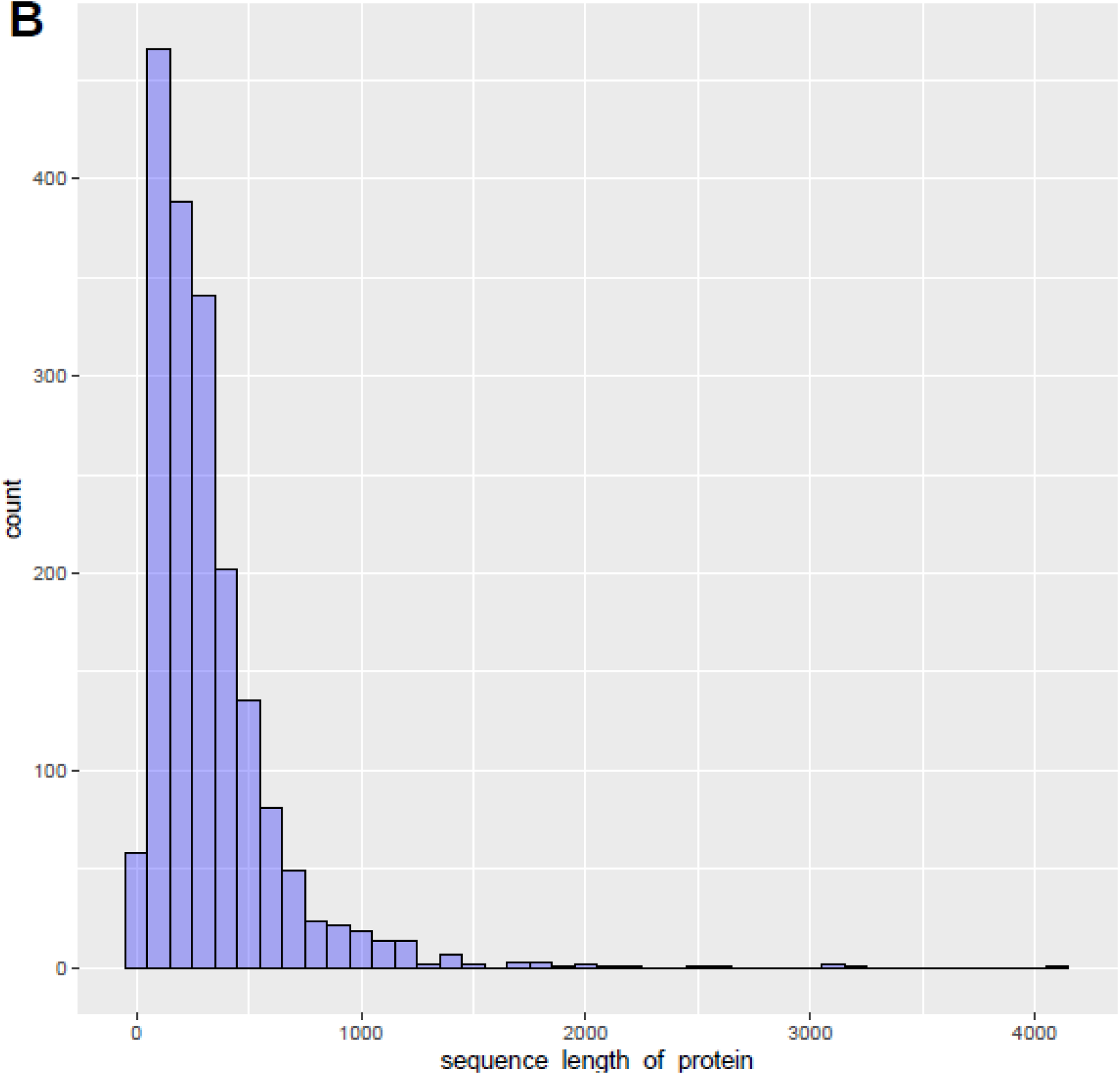
Histograms of A: number of SUMOylated lysines per target protein structure and B: sequence length of every target protein. Here count refers to frequency or number of proteins.

There are around 7618 lysines in the 1841 protein structures that conform to either the forward or reverse version of the K-x-(E/D) consensus motif. However, only 2556 or 33.5% of these lysines tend to get SUMOylated. Thus, a consensus motif alone is insufficient to guarantee SUMOylation of a lysine residue. On the other hand, all the SUMOylated lysines do not necessarily conform to the consensus motif. Majority of SUMOylated lysines analyzed in this study (4876) do not conform to the consensus motif. Thus, a sequence motif alone is not sufficient to predict all the SUMOylated lysines. Hence, the present study uses information from interactions between 3-D structures of unc9 and target proteins to predict SUMOylation sites.

The CATH database [23] classifies protein structures into different superfamilies (folds). Protein structures of the dataset used in this study were mapped to their respective CATH superfamilies (Table-7). There are 5 CATH superfamilies and members of all of these superfamilies are included in the dataset used in this study. The superfamily Alpha Beta has maximum representation as compared to other superfamilies. Information regarding subcellular localization, cellular functions and biological activity of the proteins used in this study was obtained with the help of Gene Ontology terms. There are 3 kinds of Gene Ontology terms – cellular component, molecular function and biological process. All the proteins from the dataset were mapped to their respective Gene Ontology terms and these terms were sorted in a descending order of their abundance (Tables-8 to 10).

**Table-7:** Representatives of all the CATH superfamilies are included in this study

| **CATH superfamily** | **Count** |
| --- | --- |
| Mainly Alpha | 478 |
| Mainly Beta | 338 |
| Alpha Beta | 947 |
| Few Secondary Structures | 13 |
| Special | 20 |

SUMOylated proteins mostly localize to nucleus. However, there are a few proteins that localize to cytoplasmic organelles or plasma membranes too (Table-8). Majority of the SUMOylated proteins bind nucleic acids such as DNA / RNA / ATP (Table-9). SUMOylated proteins such as zinc finger proteins are also known to bind metal ions. Biological processes such as transcription regulation, cell division, signal transduction and DNA repair have been linked to SUMOylation (Table-10).

**Table-8:** Top 10 most abundant cellular component terms

| **Rank** | **Gene Ontology cellular component terms** | **Count (Proportion %)** |
| --- | --- | --- |
| 1 | Nucleus | 419 (8.5) |
| 2 | Nucleoplasm | 411 (8.3) |
| 3 | Cytosol | 393 (7.94) |
| 4 | Cytoplasm | 325 (6.57) |
| 5 | Extracellular exosome | 179 (3.62) |
| 6 | Membrane | 127 (2.57) |
| 7 | Plasma membrane | 98 (1.98) |
| 8 | Nucleolus | 80 (1.62) |
| 9 | Mitochondrion | 70 (1.41) |
| 10 | Protein-containing complex | 68 (1.37) |

**Table-9:** Top 10 most abundant molecular function terms

| Rank | Gene Ontology molecular function terms | Count (Proportion %) |
| --- | --- | --- |
| 1 | RNA binding | 151 (3.88) |
| 2 | Identical protein binding | 148 (3.8) |
| 3 | Metal ion binding | 102 (2.62) |
| 4 | ATP binding | 101 (2.6) |
| 5 | DNA binding | 91 (2.34) |
| 6 | Zinc ion binding | 66 (1.7) |
| 7 | Chromatin binding | 62 (1.59) |
| 8 | Protein homodimerization activity | 56 (1.41) |
| 9 | RNA polymerase II cis-regulatory region sequence-specific DNA binding | 53 (1.36) |
| 10 | Enzyme binding | 50 (1.29) |

**Table-10:**
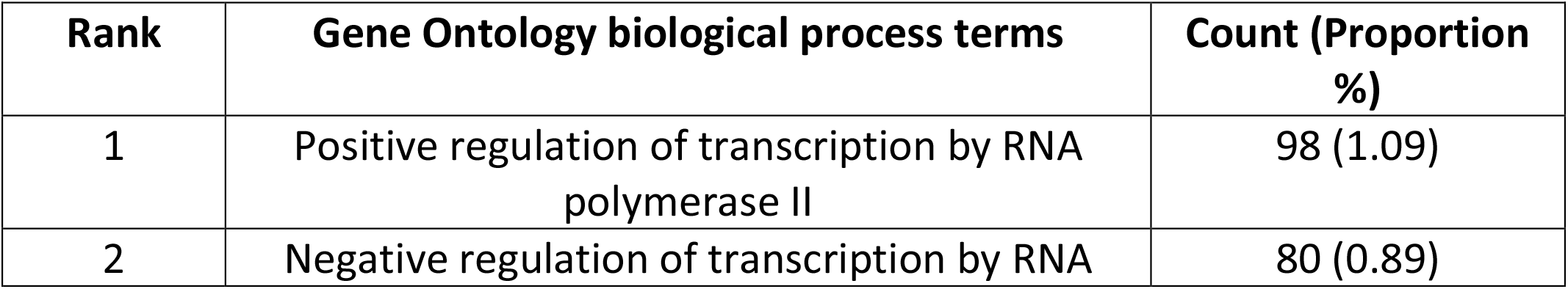
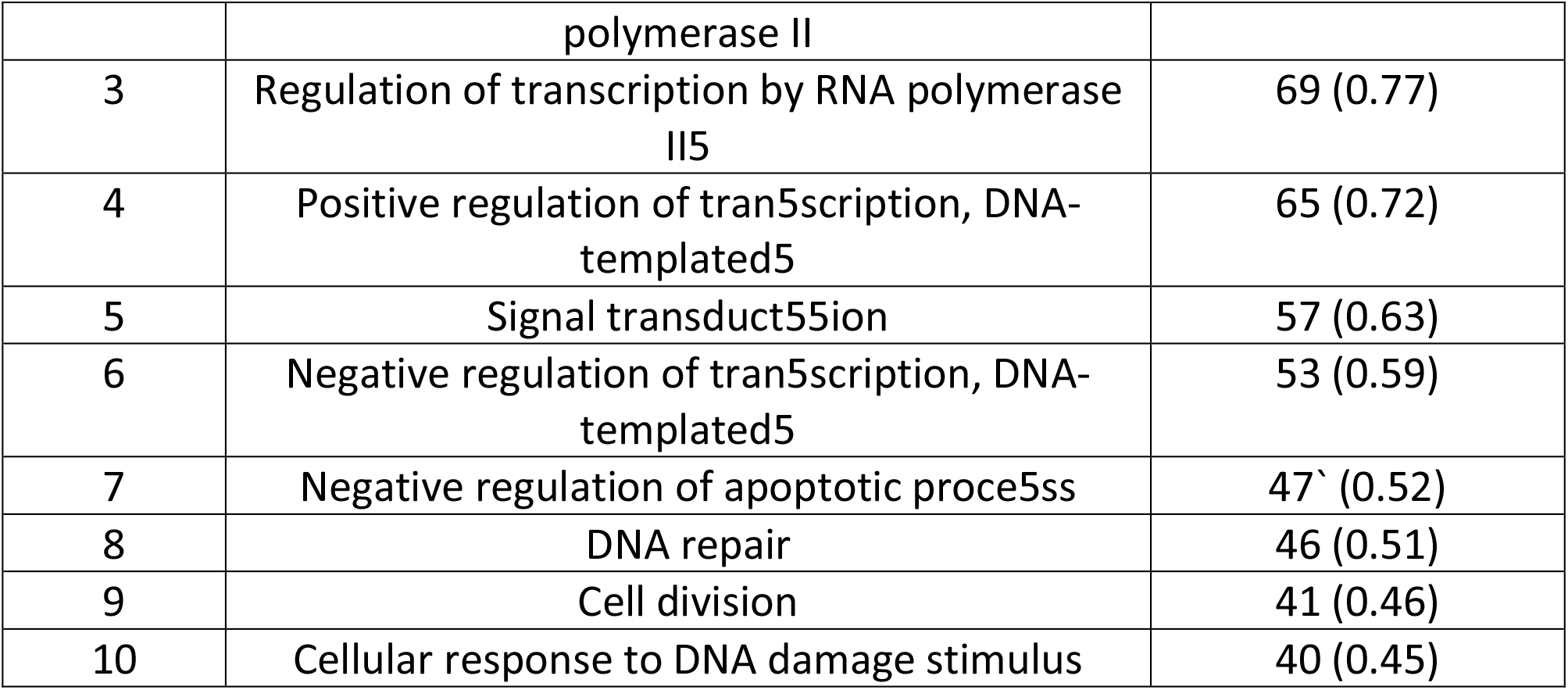
Top 10 most abundant biological process terms

### 3.2 Predictions made using residue contacts

The sampling method was applied to every lysine in all the target proteins. Clash-free poses were obtained for around half of all the SUMOylated and non-SUMOylated lysines (Table-11). For the remaining lysines, clash-free poses could not be obtained because either the main chain atoms of the target protein and ubc9 had collisions. This could be either due to unfavorable phi and psi angles of the lysine residues or the target protein conformation may not be optimal for binding ubc9.

**Table-11:** Overview of clash-free poses generated by the sampling method

|  | **SUMOylated lysines**<br>**(7432)** | **Non-SUMOylated lysines**<br>**(27874)** |
| --- | --- | --- |
| Lysines with clash-free poses | 4006 | 13022 |
| Lysines without clash-free poses | 3426 | 14852 |

A vast majority of lysines with clash-free poses do not conform to consensus motif (Table-12). There are 613 SUMOylated and 1949 non-SUMOylated lysines that have poses wherein a E/D binds positively charged patch on ubc9. But the E/D residue is not at +2 / -2 position with respect to the sequential position of lysine of interest. These E/D residues are referred to as non-consensus.

**Table-12:** Proportion of lysines with clash-free poses that conform to consensus motif

|  | **K-x-(E/D)** | **(E/D)-x-K** | **Non-consensus**<br>**(E/D)** |
| --- | --- | --- | --- |
| Clash-free SUMOylated lysines (4006) | 42 | 29 | 613 |
| Clash-free non-SUMOylated lysines (13022) | 51 | 36 | 1949 |

The Apriori algorithm generated res-pair sets varying in size from 1 to 18. Predictions were made independently for every res-pair set according to their size. Thus, predictions were made for all size-1 sets, size-2 sets and so on. The best predictions in terms of MCC were obtained for size-3 res-pair sets (Table-12). Predictions for size-3 sets were made by varying the cutoff from 1 to 185. Here cutoff refers to the number of size-3 res-pair set present in a given conformational pose. All poses having more res-pair sets than the cutoff were chosen as positive predictions or else they were marked as negative prediction.

The prediction method described here achieved a sensitivity = 27%, specificity = 98%, accuracy = 81% and MCC = 0.4 (Table-13). Our method has higher specificity than sensitivity. This can be attributed to the higher number of non-SUMOylated poses (13022) than SUMOylated poses (4006). Future prediction tools can overcome this issue by under-sampling non-SUMOylated poses or over-sampling SUMOylated poses.

**Table-13:** Overview of predictions made for size-3 res-pair sets

| Statistical parameter | Value |
| --- | --- |
| True positives | 1091 |
| True negatives | 12763 |
| False positives | 259 |
| False negatives | 2915 |
| Sensitivity | 0.272 |
| Specificity | 0.98 |
| Accuracy | 0.814 |
| F1 score | 0.407 |
| MCC | 0.396 |

Majority of lysines predicted using size-3 res-pair sets do not conform to consensus motif (Table-14). This trend is similar to the trend observed for lysines with clash-free poses (Table-12). An interesting observation is that true positives have more consensus lysines than false positives (20 versus 1).

**Table-14:**
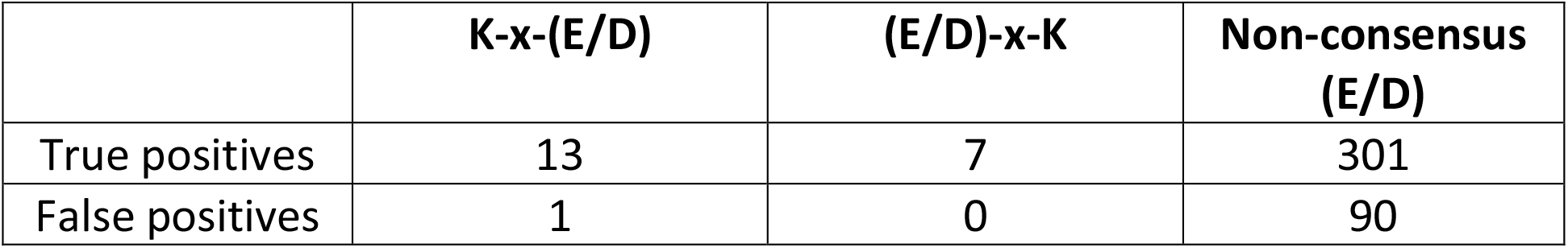
Proportion of predicted lysines that conform to consensus motif

|  | K-x-(E/D) | (E/D)-x-K | Non-consensus (E/D) |
| --- | --- | --- | --- |
| True positives | 13 | 7 | 301 |
| False positives | 1 | 0 | 90 |

There were 7826 size-3 res-pair sets that showed positive enrichment in SUMOylated poses than non-SUMOylated poses. Out of these, top 10 res-pair sets in terms of their enrichment values are given above (Table-15). There are 2 sets of particular interest. These are 87;MET, 91;ASP, 74;ASP and 76:ASP, 87;TRP, 89;ASP. Both these sets represent contacts between an aspartate residue in target protein and positively charged patch on ubc9.

**Table-15:** Top 10 size-3 res-pair sets that show maximum enrichments

| Res-pair set | Occurrence in true positives | Occurrence in false positives |
| --- | --- | --- |
| 74;LEU, 88;LEU, 91;LEU | 8 | 2 |
| 131;TYR, 129;TYR, 135;TYR | 7 | 5 |
| 139;GLU, 136;ARG, 133;ARG | 7 | 6 |
| 98;ALA, 99;GLN, 98;GLN | 6 | 2 |
| 72;ARG, 99;ARG, 65;ASN | 6 | 1 |
| 99;ARG, 71;ARG, 76;GLU | 6 | 3 |
| 87;MET, 91;ASP, 74;ASP | 6 | 4 |
| 76;ASP, 87;TRP, 89;ASP | 5 | 1 |
| 131;SER, 129;TYR, 88;TYR | 5 | 1 |
| 88;LEU, 129;LEU, 91;LEU | 5 | 4 |

Secondary structure environment of SUMOylated lysines was determined using write_data() function in MODELLER. SUMOylation targets lysine residues in all secondary structures such as alpha helices, beta sheets or coils (Table-16). The sampling method and res-pair based predictions were able to detect SUMOylated lysines irrespective of their secondary structures (Table-16).

**Table-16:** Overview of secondary structures of SUMOylated lysines

| Secondary structure | All SUMOylated lysines in the dataset (7432) | SUMOylated lysines with clash-free poses (4006) | True positives (1091) |
| --- | --- | --- | --- |
| Beta strand | 1300 | 481 | 155 |
| Loop | 3095 | 1845 | 422 |
| Alpha helix | 2988 | 1658 | 504 |
| Kink | 49 | 22 | 10 |

SUMOylated lysines that follow a consensus motif occur in all kinds of secondary structures (Table-17). Sampling and prediction methods preferred lysines in a coil / loop. This is in agreement with lysine 524 in RanGAP1, which also happens to be in a located in a loop.

**Table-17:** Secondary structures of SUMOylated lysines in consensus motif

| **Secondary structure** | **All SUMOylated consensus motif lysines (2556)** | **SUMOylated consensus motif lysines with clash-free poses (71)** | **Consensus motif lysines in True positives (20)** |
| --- | --- | --- | --- |
| Beta strand | 443 | 15 | 4 |
| Loop | 1087 | 50 | 16 |
| Alpha helix | 1011 | 6 | 0 |
| Kink | 15 | 0 | 0 |

## 4 Discussion

SUMOylation targets proteins varying in size from 100 amino acids to more than 10000 amino acids. The modification could target either one lysine in a given protein or more than 20 lysines. Proteins belonging to different folds (CATH superfamilies) are targeted by the modification. SUMOylation targets co-localize to either nucleus of a cell or to different cytoplasmic organelles and even the plasma membrane. The target proteins could possibly bind DNA / RNA / ATP and might be involved in regulation of transcription activity, cell division, DNA repair or signal transduction. SUMOylated lysines can occur in any of the 3 secondary structure environments such as alpha helices, beta sheets or loops / coils.

Almost all of the proteins used in this study were not bound to ubc9. Hence, the conformation used for sampling method may not necessarily be optimal for binding ubc9. Apart from this, factors such as crystal contacts could also influence protein conformations [24].Hence, the poses generated by the sampling method have to be analyzed with caution. Sampling different conformational poses and scoring those poses are two important aspects of any general protein-protein docking tool. In the present study, those two concepts were used for studying ubc9-target protein interactions. At present, the Protein Data Bank has structural information for only one ubc9-target complex. As more structural information becomes available for these interactions, more robust structure based prediction tools can be developed. In cases of SUMOylated proteins with unknown 3-D structures, information from Alphafold models could be used [25]. In addition, future prediction tools can achieve improved accuracy by taking into account information about SUMO E3 ligases.

## 6. Supplementary Data

All the raw data relevant to the present study can be found on Github here - https://github.com/yogendra-bioinfo/structure-based-SUMOylation-prediction. The significance of each data file can be found in a file named README.txt.

